# *In vitro* characterization of the full-length human dynein-1 cargo adaptor BicD2

**DOI:** 10.1101/2022.01.05.474964

**Authors:** Robert Fagiewicz, Corinne Crucifix, Torben Klos, Célia Deville, Bruno Kieffer, Yves Nominé, Johan Busselez, Paola Rossolillo, Helgo Schmidt

**Affiliations:** Institut de Génétique et de Biologie Moléculaire et Cellulaire, Integrated Structural Biology Department, Illkirch, France; Centre National de la Recherche Scientifique, UMR7104, Illkirch, France; Institut National de la Santé et de la Recherche Médicale, U1258, Illkirch, France; Université de Strasbourg, Illkirch, France; Department of Cellular and Molecular Medicine, University of California, San Diego, La Jolla, CA 92093, USA

## Abstract

Cargo adaptors are crucial in coupling motor proteins with their respective cargos and regulatory proteins. BicD2 is a prominent example within the cargo adaptor family. BicD2 is able to recruit the microtubule motor dynein to RNA, viral particles and nuclei. The BicD2-mediated interaction between the nucleus and dynein is implicated in mitosis, interkinetic nuclear migration (INM) in radial glial progenitor cells, and neuron precursor migration during embryonic neocortex development. *In vitro* studies involving full-length cargo adaptors are difficult to perform due to the hydrophobic character, low-expression levels, and intrinsic flexibility of cargo adaptors. Here we report the recombinant production of full-length human BicD2 and confirm its biochemical activity by interaction studies with RanBP2. We also describe pH-dependent conformational changes of BicD2 using cryoEM, template-free structure predictions, and biophysical tools. Our results will help define the biochemical parameters for the *in vitro* reconstitution of higher order BicD2 protein complexes.

## INTRODUCTION

Cytoplasmic dynein drives many retrograde microtubule transport events in eukaryotic cells. The main isoform – cytoplasmic dynein-1 (hereafter dynein-1) is involved in cell division, the transport of organelles and vesicles, brain and muscle development, and can also be hijacked by pathogenic viruses to reach cellular locations (Reck-Peterson et al., 2018;,Paschal and Vallee, 1987;,Dodding and Way, 2011;,Wilson and Holzbaur, 2012;,Fu and Holzbaur, 2014). It is also a particularly interesting motor in neurons, as it carries essential signals and organelles from distal axon sites to the cell body (Schiavo et al., 2013).

Dynein-1 motility and cargo specificity depends on cargo adaptor proteins. Currently there are eleven dynein-1 cargo adaptors known and this number is continuously growing (Olenick and Holzbaur, 2019). Cargo adaptors are characterized by α-helices forming extended coiled-coil domains and they can interact with multiple proteins. Generally, the N-terminal region of the cargo adaptor associates with the dynein motor while the C-terminal domain recruits cargos. Selectivity towards their cargos is achieved by adaptor-specific cargo binding mechanisms (Schroeder et al., 2014;,Schroeder and Vale, 2016;,Gama et al., 2017).

The best studied family of cargo adaptors are the BicD proteins. Mammals have two BicD orthologues, BicD1 and BicD2. BicD’s form homodimers and based on secondary structure predictions, they fold into the three coiled-coil domains CC1, CC2, and CC3 (Figure 1 A). The N-terminal CC1 coiled-coil domain was shown to bind dynactin and dynein via the ARP1 subunit of dynactin and the N-terminus of the dynein heavy chain (Chowdhury et al., 2015; Urnavicius et al., 2015) as well as LIC1 of the dynein motor complex (Schroeder et al., 2014; Lee et al., 2018). These multiple interactions increase the affinity of dynein for dynactin and are required for stable dynein/dynactin/BicD2 complex (DDB) formation (Schlager et al., 2014; Splinter et al., 2012; McKenney et al., 2014). It was also shown that the N-terminal region of BicD2 alone is able to induce DDB complex formation with higher efficiency than the full-length protein (Hoogenraad et al., 2003). The CC1 domain of BicD’s contains the dynein and dynactin interacting motifs – the CC1 box (Gama et al., 2017; Lee et al., 2018; Celestino et al., 2019) and the spindly motif (Gama et al., 2017; Zhang et al., 2017; Torisawa et al., 2014). The C-terminal domains of BicD’s, CC2 and CC3, are usually implicated in cargo recruitment. The BicD1 C-terminal CC3 domain can interact with GTP-bound RAB6B (Wanschers et al., 2007; Hoogenraad et al., 2001; Short et al., 2002; Schlager et al., 2010; Matanis et al., 2002) and cytomegalovirus/HHV-5 protein UL32 (Indran et al., 2010). BicD2 is able to recruit RAB6A (Peeters et al., 2013), NEK8 (Holland et al., 2002), DNAI1 (Oates et al., 2013; Peeters et al., 2013), and kinesin KIF5A (Splinter et al., 2010). BicD2 C-terminus is also able to interact with RANBP2 (Splinter et al., 2010) (or NUP358), a component of the nuclear pore complex. The subsequent BicD2 mediated recruitment of dynein-dynactin (Figure 1B) allows the tethering of the centrosome to the nucleus prior to mitosis (Splinter et al., 2010) and drives interkinetic nuclear migration during the development of the human neocortex (Hu et al., 2013; Baffet et al., 2015). It has been shown in vivo and in vitro that BicD2 is recruited to RanBP2 in a CDK1-dependent manner (Baffet et al., 2015) (Figure 1B).

**Figure 1.**
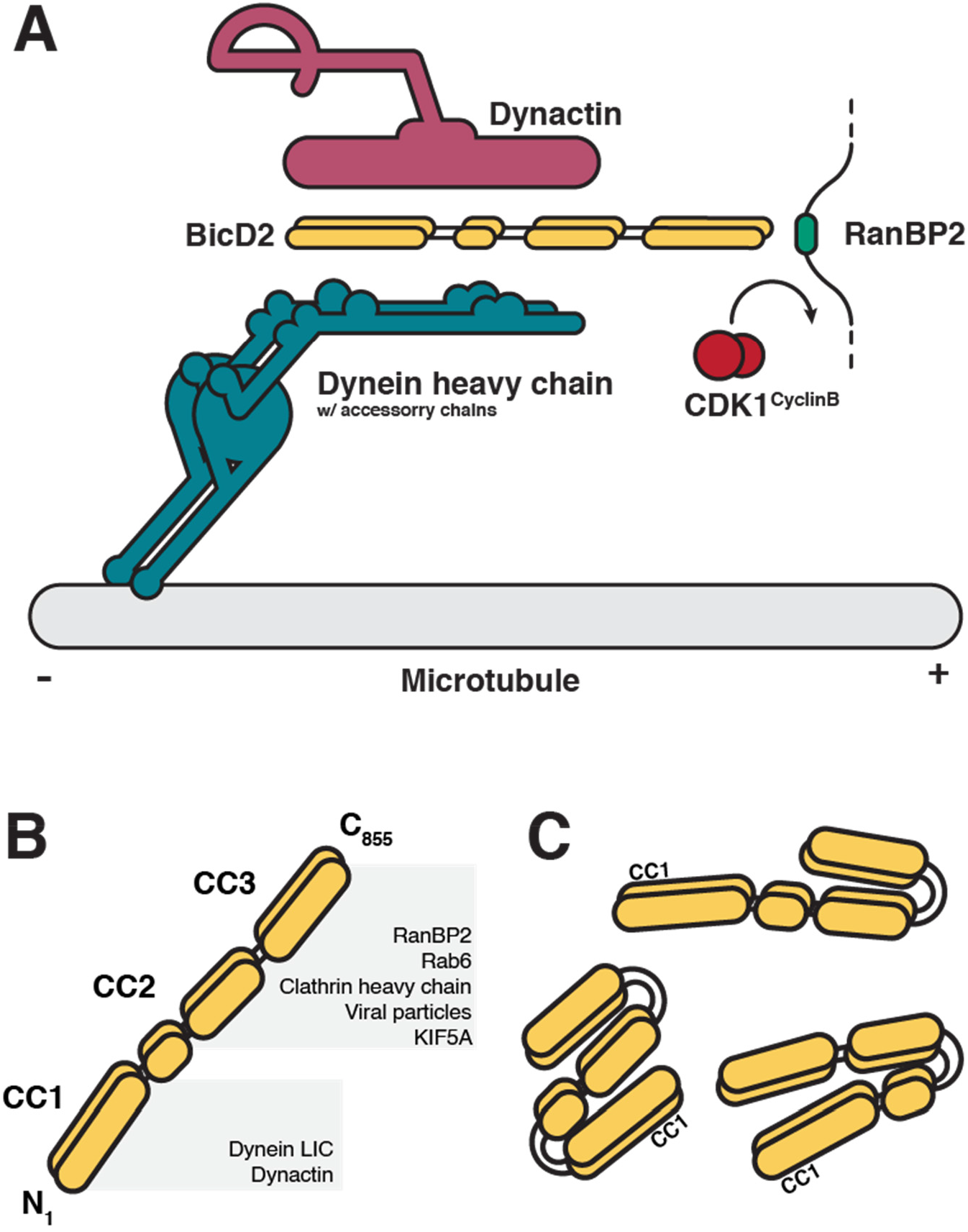
Schematic representation of BicD2 domain organization and function. (A) Domain organization of the BicD2 cargo adaptor dimer. The N-terminal part is responsible for binding dynein and dynactin while the C-terminal part binds the plus-end directed motor kinesin-1 and cargos like RanBP2. (B) BicD2 associates with its RanBP2 nuclear pore cargo and dynein-dynactin. The CDK1^CyclinB^ mediated phosphorylation of RanBP2 triggers the binding of BicD2 to recruit dynein-dynactin to the nucleus during interkinetic nuclear migration in the developing neocortex. (C) Hypothetical BicD2 autoinhibited conformations from the previously reported data.

Cargo adaptors of the BicD family also interact with themselves. It has been observed by several groups that the C-terminal part is able to make contact with the N-terminal part leading to an autoinhibited state in BicD1 (Terawaki et al., 2015) and BicD (Liu et al., 2013; Wharton and Struhl, 1989) (Figure 1C). There is little known about human BicD2 autoinhibition and many features are assumed to be common with the better-studied *Drosophila* BicD ortholog (Sladewski et al., 2018; Stuurman et al., 1999). It has also been suggested that the autoinhibited state can be relieved by cargo binding (Liu et al., 2013; Huynh and Vale, 2017; McClintock et al., 2018; Sladewski et al., 2018). There is evidence of cargo-induced autoinhibition release in *Drosophila* BicD with Egl and mRNA as a cargo (Sladewski et al., 2018). So far no such studies have been done for human full-length BicD1 or BicD2.

Understanding the structure, dynamics and reactivity of cargo adaptor proteins is required to advance our understanding of cargo selection, recruitment, and release. Most of the *in vitro* reconstitution experiments of the human BicD2 with cargo and/or dynein-dynactin were performed on a truncated protein or the *Drosophila* ortholog. As it is an extensively studied dynein adaptor, there is a need for a protocol for robust production of the full-length human BicD2. The long-standing challenges in characterising cargo adaptors like BicD2 are limited expression levels and solubility which makes them poor targets for structural and functional analyses. Here we describe the recombinant production of human full-length BicD2 and provide novel insights to its *in vitro* behaviour. We validate the dimeric nature and correct folding of our recombinant BicD2. Our results also reveal unexpected pH-dependent conformational changes of BicD2 in the pH range from 6 to 8. We also address the biochemical activity of BicD2 by interaction studies with RanBP2. Our findings lay the ground for further studies on BicD2 reactivity, regulation, and role in motor protein complexes formation.

## RESULTS

### Biochemical characterization of full length BicD2

We succeeded in producing full-length human BicD2 isoform 2 under mild solubilizing conditions (see STAR Methods) and subsequently characterized its biochemical and functional properties. The addition of mild zwitterionic surfactants like CHAPS to the lysis buffer was crucial for obtaining sufficient protein amounts for large scale purifications. Our size-exclusion multi-angle Light scattering (SEC-MALS) analysis indicates a molecular weight of 192 (±0.7%) kDa (Figure S1A), which is slightly larger than the expected value of 186 kDa for a full-length BicD2 dimer The higher molecular weight is likely caused by the presence of residual CHAPS detergent. BicD2 also elutes at a volume of 1.35 mL on a Superose^®^ 6 Increase 3.2/300 size exclusion column, which is earlier than expected and most likely caused by the increased hydrodynamic radius due to the extended coiled-coil fold of BicD2.

We evaluated the effect of buffers on the stability of BicD2 by differential scanning fluorimetry (nanoDSF) to establish an optimal buffer composition. The pH screen revealed that the BicD2 stability varies considerably in the range from pH 6.0 to pH 8.0 (Figure 2A). The melting curves at pH 7.0 and pH 8.0 show clear but distinct inflection points, while the curve at pH 6 indicates partly unfolded protein. Full-length BicD2 carries a net negative charge at all these pH values (Figure S1B). We also performed three SEC runs on full-length BicD2 at pH 6.0, 7.0, and 8.0. The first run after affinity purification at pH 7.0 and pH 8.0 was characterized by an elution peak for dimeric BicD2 between 1.3 – 1.4 mL and a broad peak of degraded BicD2 around 1.8 mL (Figure S1C). The second SEC run yielded clean BicD2 fractions (Figure 2B). The slightly different SEC elution volumes and the distinct inflection points in the DSC melting curves suggest the possibility of different BicD2 conformations at pH 7.0 and pH 8.0. Moreover, the SEC run at pH 7.0 had a significantly higher ratio of full length BicD2 to degraded BicD2 compared to the SEC run at pH 8.0, where degradation proceeds more rapidly (Figure S1C). At pH 6.0 BicD2 eluted almost at the end of the column suggesting partial denaturation (Figure 2B). Re-running the BicD2 peak fractions revealed elution peaks at the expected volumes without additional degradation peaks. These results were further confirmed by circular dichroism (CD) experiments (Figure S1D). The spectra for pH 7.0 and pH 8.0 suggest almost pure α-helical content, consistent with the predicted BicD2 fold. The CD spectrum for pH 6.0 does not resemble the α-helical model, but also does not fit into random-coil models. Heating the protein up to 60 °C did not reveal the random coil characteristic peaks (Figure S1E). This suggests that the protein at pH 6.0 adopts a partially unfolded structure.

**Figure 2.**
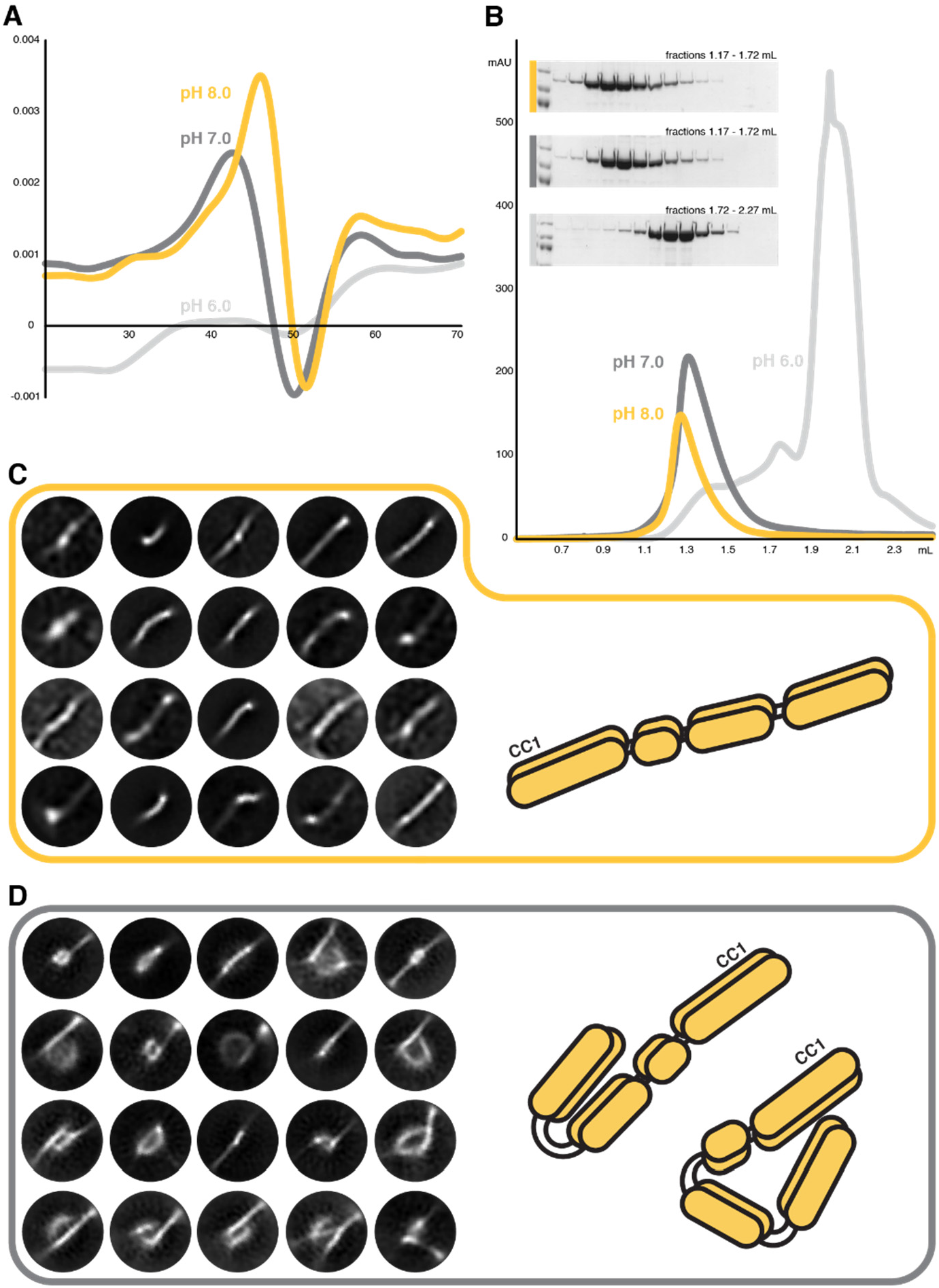
Biochemical and structural characterization of human full-length BicD2 at different pH values. (A) First derivative of the DSF melting curves at three different pH values. The curves at pH 7.0 and pH 8.0 display main inflection point in the range of 47-50°C. The curve at pH 6.0 does not show a distinct inflection point. Each measurement was repeated 3 times. (B) SEC profiles of BicD2 run at three different pH values. Each sample was run twice on a Superose^®^ 6 Increase column to ensure clean peak fractions. (C) and (D) Representative cryoEM 2D classes from ~700 micrographs (per pH condition). The yellow box represents classes from pH 8.0 while the grey box from pH 7.0. In both cases the data was processed with Relion3.1. See also Figures S1 and S4.

### Influence of pH value on BicD2’s conformation and stability

In order to obtain insights into the conformation of BicD2 at pH 7.0 and pH 8.0, we directly visualized BicD2 at these pH values using a cryo-electron microscope. We collected two datasets and applied identical data processing workflows to avoid bias in the interpretation. The representative 2D classes showed that most molecules at pH 8.0 fall into an extended conformation (Figure 2C), whereas the vast majority of BicD2 particles adopt more compact conformations (Figure 2D) at pH 7.0. These results are consistent with the SEC runs where BicD2 elutes at 1.4 mL at pH 7.0 but 1.35 mL at pH 8.0 suggesting a shorter hydrodynamic radius at pH 7.0.

Next, we wanted to identify amino-acid residues that potentially could be responsible for the pH dependent conformational changes of BicD2. We analysed multiple sequence alignments and focused on conserved histidines as they often have crucial functions in pH-sensitive proteins. An alignment of BicD, BicD1, and BicD2 sequences from all animal kingdoms showed only a poor sequence identity of less than 10%. However, in the C-terminal part of the CC2 domain there is a conserved ‘YH’ sequence motif and an additionally conserved histidine (H640 in human isoforms) that is present in all vertebrates (Figure 3A). We generated a structure prediction with AlphaFold2 (Jumper et al., 2021) (AF2) for the CC2 and CC3 domains of human BicD2 (Figure 3B) to evaluate where these residues are potentially located within the predicted structure. Our analysis suggests that they are not involved in dimer interactions, but that they are located within the tri-helical interface of H3, H4, and H5. We compared the YH-containing motif of *hs*BicD2 (Y538-H539) with its two distant isoforms – *D. melanogaster* BicD and *C. elegans* BicD1. The architecture is very similar for all isoforms. In the case of hsBicD2, the additionally conserved histidine residue H640 on helix H5 could interact with residue Y538 on helix H3 (Figure 3C). In order to validate the structure predicted by AF2, we have also compared it to the negative stain class averages reported for *D. melanogaster* BicD (Sladewski et al., 2018) (Figure S2A, B and C).

**Figure 3.**
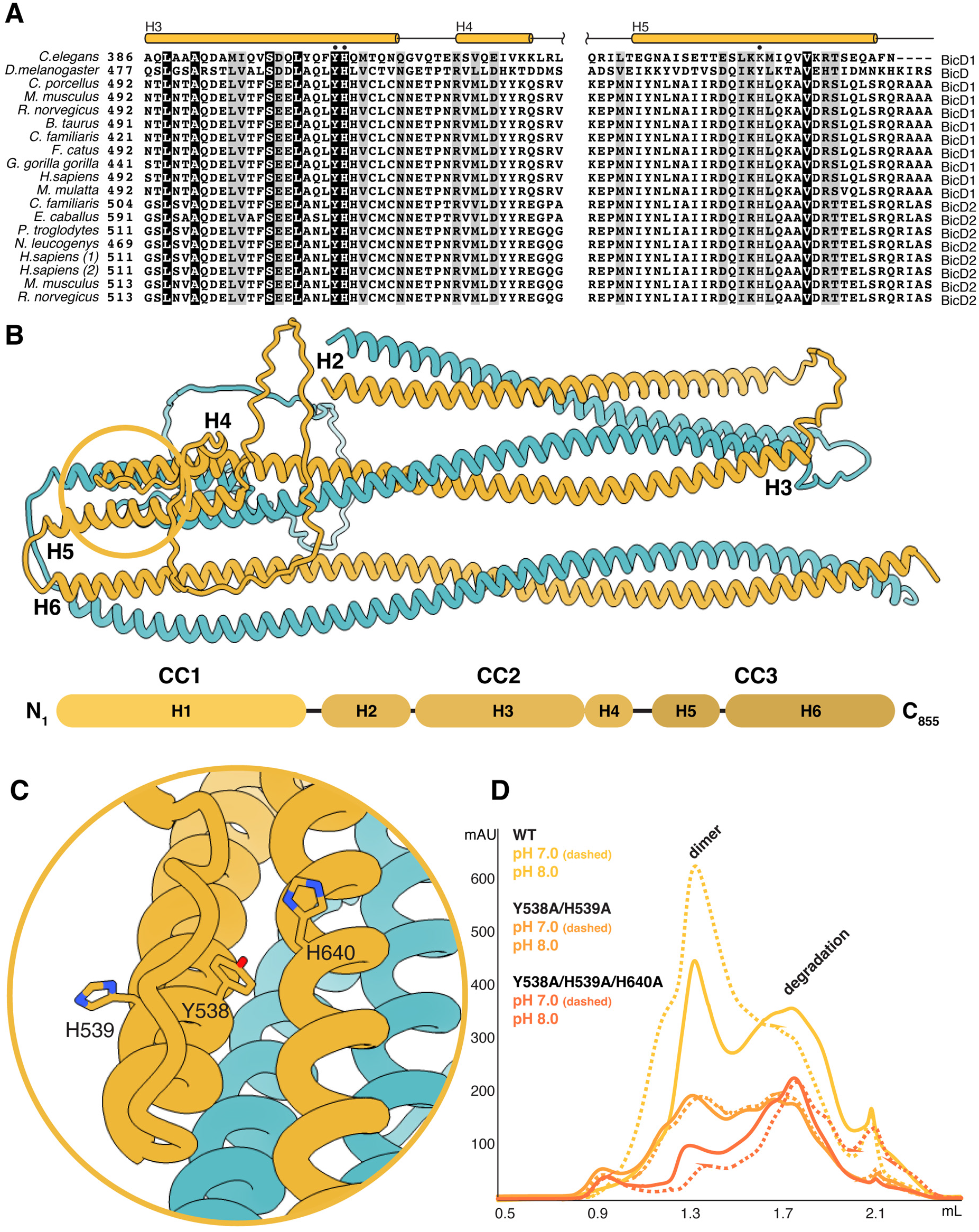
The conserved amino-acid residues Y538,H539 and H640 contribute to BicD2 stability. (A) Sequence alignment of the tri-helical domain (H3, H4, and H5 helix) of *D. melanogaster, C. elegans,* and mammalian BicD1 and BicD2 isoforms. The conserved amino-acid residues Y538, H539 and H640 are marked by black dots. (B) Structure of the BicD2 CC2 and CC3 dimer predicted by AlphaFold2. The C-terminal loop of CC3 has been deleted for clarity. (C) Enlarged view of the conserved histidine and tyrosine residues that are crucial for protein stability. (D) Chromatograms of affinity purified BicD2 variants. Yellow - WT, orange - Y538A/H539A double mutant, dark orange - Y538A/H539A/H640A triple mutant. In all chromatograms the solid line represents pH8.0, and the dashed line pH 7.0. For each construct an identical protein amount was loaded. See also Figure S2.

To test if the YH motif and the interhelical tyrosine-histidine pair (Y538-H640) interfere with BicD2’s pH dependent behaviour *in-vitro,* we generated the H539A/Y538A and H539A/Y538A/H640A mutants and characterized them by SEC at pH 7.0 and pH 8.0. The fact that the differences in the BicD2 elution peaks do still exist suggests that also the underlying conformational differences are still present. Furthermore, the mutant BicD2 proteins are more prone to degradation because the equilibrium between the peaks for full-length BicD2 and its degradation products has shifted towards the degradation products. This effect is more pronounced for the H539A/Y538A/H640A triple mutant (Figure 3D). At pH 7.0 the BicD2 fraction is the most abundant, while at pH8.0 the BicD2 fraction significantly decreases, and the protein is prone to degradation. Even though H539/Y538/H640 does not seem to be involved in the pH-induced conformational changes of full-length BicD2, these residues nevertheless are clearly important for the stability of the protein.

### BicD2’s interaction with RanBP2

RanBP2 (or Nup358) is known to recruit BicD2 through a small disordered region between two Ran binding domains (2003-2444) as demonstrated in yeast two-hybrid assays (Splinter et al., 2010). The BicD2 binding domain of RanBP2 (hereafter RanBP2_BBD_) interacts with the C-terminus of BicD2 (715-804) (Gibson et al., 2021) and contains several consensus CDK1 phosphorylation sites, which have been shown to be important for BicD2 binding (Baffet et al., 2015). The individual contribution of these phosphorylation sites to BicD2 binding is unknown. A recent *in-vitro* experiment demonstrated that the RanBP2_BBD_-BicD2 complex can also be formed in the absence of phosphorylation (Gibson et al., 2021). Since all these experiments used the truncated BicD2 C-terminus, the binding of the full-length BicD2 to RanBP2_BBD_ remains to be characterized. In order to clarify the role of phosphorylation in BicD2-RanBP2_BBD_ complex formation and to confirm the biochemical activity of our recombinant full-length BicD2, we evaluated the interaction with its binding partner RanBP2.

We performed double pull-down assays with full length BicD2 and RanBP2_BBD_ as well as with a phosphomimetic variant of RanBP2_BBD_ where the known CDK1 phosphorylation sites had been mutated to aspartates (T2153D, S2246D, S2251D, S2276D, and S2280D). We were able to obtain a BicD2-RanBP2_BBD_ complex in both cases (Figure S3A). However, subsequent SEC runs of the purified complexes indicated that they are not stable and that they dissociate. In order to investigate if RanBP2_BBD_ phosphorylation leads to more stable complex formation, we produced recombinant CDK1 (Veld et al., 2021) and performed an *in vitro* kinase assay on RanBP2_BBD_. The mass spectrometry analysis revealed eight S/T consensus sites in the BicD2 binding region of RanBP2 of which five overlapped with earlier findings (Baffet et al., 2015) (Figure S3B). We mixed *in vitro* phosphorylated or non-phosphorylated RanBP2_BBD_ with full length BicD2 and analysed the samples by SEC. Successful complex formation was only observed for *in vitro* phosphorylated RanBP2 (Figure 4A). Next, we investigated if BicD2 can be phosphorylated by CDK1 and if such a modification could potentially strengthen complex formation with RanBP2_BBD_. The mass spectrometry analysis after BicD2 *in vitro* phosphorylation indicated five phosphorylation sites in regions predicted to be structurally disordered demonstrating that BicD2 can be phosphorylated by CDK1 (Figure S3B). We mixed the *in vitro* phosphorylated BicD2 with non-modified RanBP2_BBD_ and subsequently analysed the sample by SEC. There was no indication of complex formation (Figure 4A). The equivalent experiment with *in vitro* phosphorylated RanBP2_BBD_ did not show any evidence of increased BicD2-RanBP2_BBD_ complex formation compared to non-modified BicD2 – *in vitro* phosphorylated RanBP2 SEC run (Figure 4A). These results suggest that BicD2 phosphorylation does not contribute to the BicD2-RanBP2_BBD_ complex formation.

**Figure 4.**
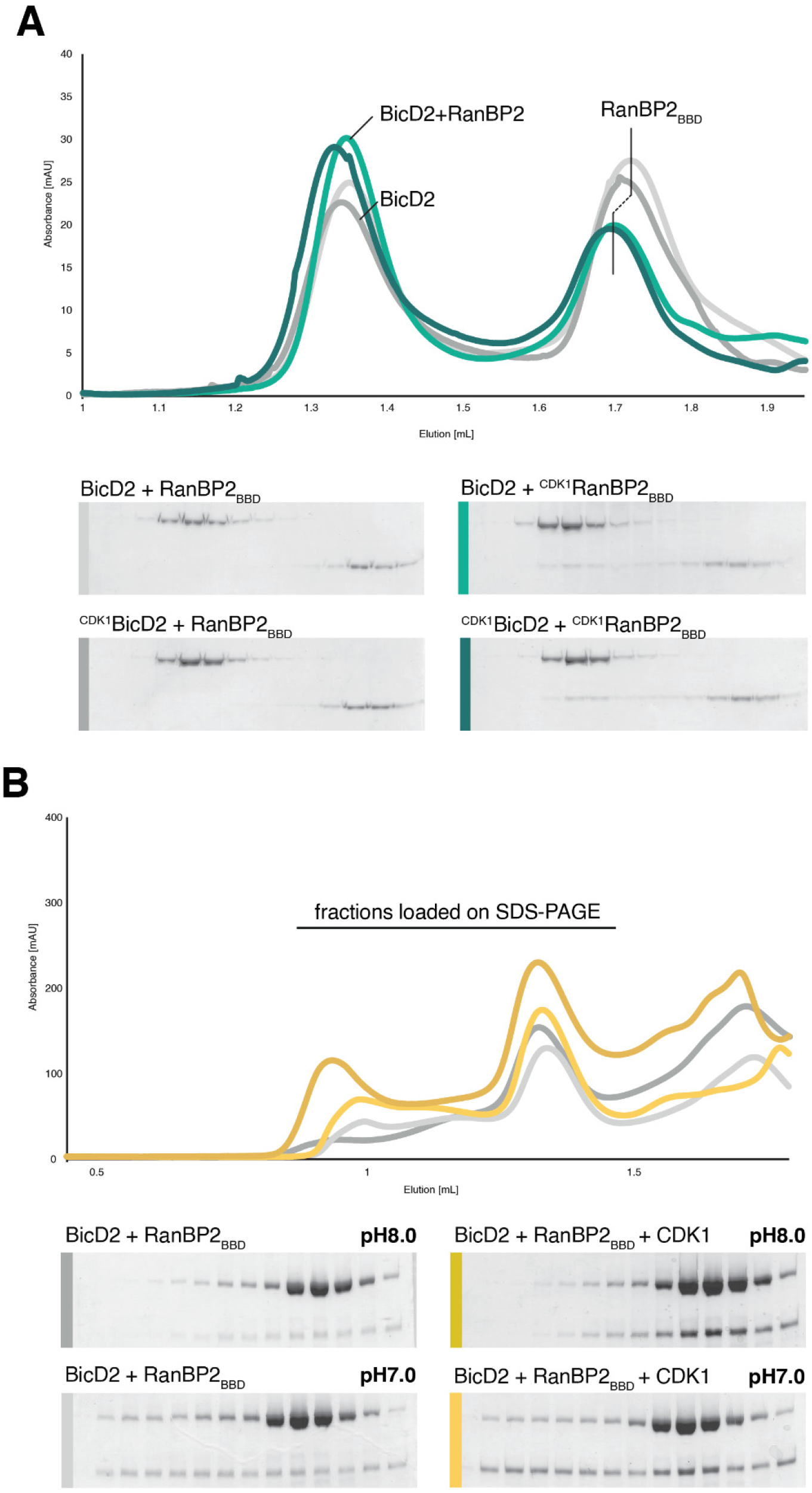
SEC profiles of the BicD2 reaction with RanBP2_BBD_ in the absence and presence of CDK1^CyclinB^. (A) Purified proteins were mixed at a 1:1 molar ratio and incubated for 30 minutes at RT. In light grey a chromatogram of unphosphorylated BicD2 and unphosphorylated RanBP2_BBD_. In dark grey CDK1^CyclinB^ phosphorylated BicD2 with unphosphorylated RanBP2. In light green unphosphorylated BicD2 with CDK1^CyclinB^ phosphorylated RanBP2, and dark green represents both proteins being CDK1^CyclinB^ phosphorylated. All reactions were performed at pH 8.0. (B) Pull-down of affinity purified BicD2, RanBP2_BBD_, at two different pH values with or without CDK1^CyclinB^. The dark yellow chromatogram represents the BicD2, RanBP2_BBD_, and CDK1^CyclinB^ pull-down at pH 8.0, while light yellow shows the same reaction but at pH 7.0. Dark and light grey chromatogram show the same reactions but without activity of CDK1^CyclinB^ complex (at pH 8.0 and pH 7.0 respectively). See also Figures S3 and S4.

We also analysed the influence of pH on BicD2-RanBP2_BBD_ complex formation. To this end, we mixed full-length BicD2, RanBP2_BBD_ and CDK1 at pH 7.0 and pH 8.0 to induce complex formation and analysed the samples by SEC. We also carried out the corresponding control experiments in the absence of CDK1 (Figure 4B). From the fractions analysed by SDS-PAGE, it is evident that complex formation is enhanced with CDK1-treated samples. Interestingly, the two runs at pH 7.0 (with and without CDK1) showed a broad distribution of RanBP2_BBD_ throughout the elution and only minor increase in the RanBP2 intensity at BicD2 peak fractions in the presence of CDK1. Taken together, these observations suggest that the BicD2-RanBP2_BBD_ complex forms best under slightly alkaline pH which narrows the pool of RanBP2_BBD_ species.

## DISCUSSION

The ability of Dynein-1 to bind a wide range of cargoes is mediated by its compatibility with many cargo adaptor proteins, such as BicDs, Hooks, NINs, Spindly, TRAKs, RILP or JIP3 (Olenick and Holzbaur, 2019). Investigating *in vitro* reconstituted complexes between cargo adaptors and dynein/dynactin has led to valuable insights into dynein motor recruitment and activation (Terawaki et al., 2015; Liu et al., 2013; Noell et al., 2019). However most of these studies were done with truncated BicD2 constructs that lack either the dynein or the cargo -binding domains. Many biophysical and structural studies have been hampered by the lack of sufficient amounts of full-length cargo adaptors. Here we report a robust method for the recombinant production of a full-length BicD2 and demonstrate its correct fold and biochemical activity towards its RanBP2 cargo. We believe that other full-length cargo adaptors might benefit from the expression and purification protocols described here and that our results will facilitate *in vitro* reconstitution approaches of motor-cargo complexes.

One of the unexpected findings was the strong influence of the pH value on the conformation and stability of full-length BicD2. At pH 6.0, 7.0 and 8.0 BicD2 switches from partially unfolded to compact and to an open conformation. Our cryoEM 2D classification of full-length BicD2 at pH 7.0 indicates that over 50% of the particles fall into compact classes while the majority of particles at pH 8.0 adopt an open conformation. Recent work has provided evidence that the autoinhibited state is characterized by an interaction between the CC2 and CC3 domains (Sladewski et al., 2018). Although we were not able to assign individual coiled-coil domains in our cryoEM 2D class averages due to the intrinsic flexibility of BicD2, there is the possibility that the more compact 2D class averages at pH 7.0 might represent the autoinhibited state and the extended conformation at pH 8.0 the open, activated state. A similar autoinhibited conformation has also been recently described for cargo adaptor Spindly (d’Amico et al., 2022) which suggest that autoinhibition might be a common feature of cargo adaptors.

It is not clear if these pH driven conformational changes are relevant only *in vitro* or if they also have a physiological relevance. In principle, the cellular pH value could also trigger pH dependent conformational changes of the BicD2 cargo adaptor. At the cytosolic pH value of ~7.2 most BicD2 molecules would adapt the compact, potentially autoinhibited form and require additional, activating signals to recruit motor proteins and cargo. Conditions that lead to an increase in the intracellular pH, like mitosis and growth factor signalling (Gagliardi and Shain, 2013; Pouysségur et al., 1985; Chambard and Pouyssegur, 1986), could bypass the need for such signals and activate the BicD2 cargo adaptor on a globular level. Adaptor autoinhibition is not a BicD2-unique phenomenon. A recent study describes a similar behavior for the dynein-1 adaptor Spindly (d’Amico et al., 2022). Cargo adaptors alone are not extensively studied but our work and the one presented by D’Amico et al., (2022) suggests that this behavior might be common for some cargo adaptors and play a role in recruitment of partner proteins.

In the case of BicD2 one of the activating signals is believed to be the interaction with its cargos like RanBP2 (Liu et al., 2013; Sladewski et al., 2018; Huynh and Vale, 2017; McClintock et al., 2018). Furthermore, post-translational BicD2 modifications might also influence its reactivity. BicD2 contains several consensus phosphorylation sites (Holland et al., 2002) and an acetylation site in its N-terminal part (Van Damme et al., 2012).

We also have identified three conserved residues, H539/Y538/H640 that contribute to the stability of BicD2. Mutating these amino-acid residues to alanines clearly increased the tendency of BicD2 for degradation, which could be caused by auto-cleavage or residual protease amounts in the sample. Our template-free AF2 prediction suggest the possibility of Y538/H640 stabilizing the H3-H4-H5 α-helical bundle via an intramolecular hydrogen bond (Figure S2D). Disrupting it could trigger partial unfolding and subsequent protein degradation. A similar architecture (Figure S2D) was also observed in a pH-sensitive endotoxins and mutating the equivalent histidine-tyrosine pair mutation lead to protein destabilization and degradation (Seale, 2006).

We characterized the BicD2 reactivity with its known cargo protein RanBP2. There are conflicting results in the field with respect to the question whether RanBP2 phosphorylation is a requirement for the interaction with BicD2 (Baffet et al., 2015; Gibson et al., 2021). Our results now reconcile these reports, because they show that BicD2 can bind both non-phosphorylated as well as phosphorylated RanBP2_DDB_. In double pull-downs we observed that BicD2 readily interacts with RanBP2 without any protein modification at pH 7.0 and 8.0. These complexes are nevertheless unstable over SEC. This suggests that there is a weak association of these proteins *in vitro* in a wide range of conditions. The CDK1 phosphorylation of RanBP2_BBD_ (but not BicD2) significantly increased the stability of the complex and increase in the pH value improved the complex homogeneity.

## Supporting information

Key resource table

## ACKNOWLEDGEMENT

This research was supported by a LabEx start up grant (ANR-10-LABEX-30-HS) to HS and Boehringer Ingelheim Fonds as well as FRM PhD fellowships to RF. This study was further supported by the grant ANR-10-LABX-0030-INRT, a French State fund managed by the Agence Nationale de la Recherche under the frame program Investissements d’Avenir ANR-10-IDEX-0002–02. The authors acknowledge the support and the use of resources of the French Infrastructure for Integrated Structural Biology (FRISBI) ANR-10-INBS-05 and of Instruct-ERIC. We also thank the IGBMC molecular biology and virus service (Nicole Jung and Thierry Lerouge). This work of the Interdisciplinary Thematic Institute IMCBio, as part of the ITI 2021-2028 program of the University of Strasbourg, CNRS and Inserm, was supported by IdEx Unistra (ANR-10-IDEX-0002), and by SFRI-STRAT’US project (ANR 20-SFRI-0012) and EUR IMCBio (ANR-17-EURE-0023) under the framework of the French Investments for the Future Program.

## AUTHOR CONTRIBUTIONS

RF: conceptualization, data curation and analysis, validation, investigation, methodology, funding acquisition and manuscript writing. CC: methodology, investigation and data analysis. TK: methodology, investigation and data analysis. CD & BF: data analysis and discussion. YN, JB and PR: investigation and data analysis. HS: conceptualization, data analysis, supervision, funding acquisition, methodology, manuscript writing and project administration.

## STAR METHODS

### RESOURCE AVAILABILITY

#### Lead contact

Further information and requests for resources and reagents should be directed to and will be fulfilled by the lead contact, Helgo Schmidt

#### Materials availability

Plasmids generated in this study are available upon request.

#### Data and code availability

No cryoEM or x-ray crystal structures were generated in this study. This paper does not report original code. Any additional information required to re-analyze the data reported in this paper is available from the lead contact upon request.

### EXPERIMENTAL MODEL AND SUBJECT DETAILS

E. coli BL21(DE3) cell were obtained from New England Biolabs, product number C2527H. The Sf9 insect cells used in this study were obtained from Sigma-Aldrich, product number 71104-M.

### METHOD DETAILS

#### Cloning and mutagenesis

Human BicD2 isoform 2 and RanBPBBD genes were ordered from Epoch LifeScience and codon-optimized for *Escherichia coli* expression. The genes were subsequently cloned into the pNHD expression vector, a vector used for site-specific pSer incorporation (Rogerson et al., 2015), but both proteins express equally well in a commercial pMMS vector). The BicD2 plasmid was modified to introduce an N-terminal STREP-tag followed by a short ‘GSGSG’ linker. The RanBPBBD plasmid was fused with a C-terminal double TEV cleavage site, a GSGSG linker, and a 6xHis tag. The H539A/Y538A and H539A/Y538A/H640A mutants were generated by rolling circle PCR. The co-expression plasmid encoding CDK1-cyclinB for insect cell expression was a kind gift from the Andrea Musacchio lab.

#### Protein expression and purification

Plasmids for bacterial expression were transformed into *E. coli* BL21 (DE3) chemically competent cells and grown overnight on tetracycline plates. The plate was used for inoculation of the 50 mL LB medium pre-culture which was grown overnight at 37°C for further scale-up. Protein expression was induced at 25°C with 1mM isopropyl-1-thio-β-D-galactopyranoside (IPTG) and incubated for 3h at 37°C. Cells were harvested, washed with 1xPBS, and frozen. BicD2 WT and mutants were re-suspended in lysis buffer containing 1xPBS pH8.0, 10% glycerol, 7 mM CHAPS, 2 mM DTT, 2mM PMSF and cOmplete EDTA-free protease inhibitor cocktail tablets (Roche), and incubated on a roller for 1-2h at 4°C. Cells were lysed by sonication (5×30s with 30s breaks on ice) and centrifuged at 30000 rpm at 4°C for 1 hour. The supernatant was mixed with pre-equilibrated STREP-tactin 4XL beads and incubated on a roller for 30 minutes at 4°C before loading on a gravity flow column. The resin was washed with 10 CVs of the lysis buffer, 10 CVs of the wash buffer (1xPBS pH8.0, 10% glycerol, 7 mM CHAPS), and eluted with elution buffer (1xPBS pH8.0, 10% glycerol, 7 mM CHAPS, 50 mM biotin). Pooled fractions were concentrated to 2 mg/mL and snap-frozen in 100 μL aliquots for further experiments and gel filtration. RanBP2_BBD_ was purified in the same manner as BicD2 with a difference in the buffer used (20 mM Tris-HCl pH8.0 instead of PBS). Both constructs (WT) yielded around 1 mg of protein per litre of culture after the affinity purification step.

The CDK1^CyclinB^ plasmid was transformed into *E. coli* DH10MB-MCherry chemically competent cells with a heat shock at 42°C for 30s followed by 6h recovery in LB medium at 37°C with shaking. The cells were then plated on agar plates containing kanamycin (50 μg ml^-1^), gentamicin (7 μg ml^-1^), tetracycline (10 μg ml^-1^), Xgal (600 μg ml^-1^) and IPTG (40 μg ml^-1^) and positive clones were identified by blue/white selection after 24h incubation at 37°C followed by 24h at RT. The positive clones were grown in LB media containing kanamycin (50 μg ml^-1^), gentamicin (7 μg ml^-1^), and tetracycline (10 μg ml^-1^) overnight. Bacmids were purified using the isopropanol precipitation method. 2ml of Sf9 cells at 0.5×10^6^ cells per ml were transfected with 2 μg of fresh bacmid DNA and FuGene HD transfection reagent (Promega) at a ratio of 3:1 transfection reagent to DNA. After 72h the 6-well plate was analysed for mCherry fluorescence signal, and the positive conditions (V0) were directly used for further virus amplifications by adding 2mL of V0 to 50 mL of Sf9 cells at 1×10^6^ cells per mL (V_1_) and grown for 72h. The V_1_ was then used for infecting another 50 mL of Sf9 at 1×10^6^ cells/mL (V_2_). The V_2_ virus was further used to infect 500 mL of the Sf9 culture 1×10^6^ cells/mL. After 72h the cells were collected at 1000 rpm for 10 min at 4°C. The pellet was flash-frozen in liquid nitrogen and stored at −80°C until purification. The CDK1^CyclinB^ pellet was re-suspended in the lysis buffer containing 50mM Tris-HCl pH8.0, 10mM MgCl2, 10% glycerol, 0.1% NP40, 0.1 mM EDTA, 2mM DTT, 2 mM PMSF, protease inhibitors, and incubated on a roller for 1h at 4°C. The cells were lysed by sonication (2×30s with 30s breaks on ice) and centrifuged at 40000 rpm at 4°C for 1 hour. The supernatant was mixed with pre-equilibrated GST-sepharose beads for 30 minutes at RT and loaded onto a gravity flow column. The resin was washed with 10 CV of the lysis buffer, 10 CV of the wash buffer (Tris-HCl pH8.0, 150mM NaCl, 10% glycerol, 0.1mM EDTA), and eluted with elution buffer (Tris-HCl pH8.0, 150mM NaCl, 10% glycerol, 0.1mM EDTA, 25mM reduced GST). Pooled fractions were concentrated to 0.5 mg/mL and snap-frozen in 50μL aliquots for further experiments.

#### SEC-MALS

Prior to SEC-MALS analysis, the BicD2 dimer was purified on Superose^®^ 6 Increase 3.2/300 column in SEC buffer (50 mM Tris pH 7.5, 150 mM Potassium acetate, 10 mM Magnesium acetate, 7 mM CHAPS). 90μL of BicD2 sample (1mg/mL) was injected into FPLC Ettan Micro LC (formerly GE Healthcare, now Cytiva) coupled with MiniDAWN TREOS MALS detector (Wyatt Technology), Optilab T-rEX (Wyatt Technology) RI detector, and Superdex^®^ S200 10/300 GL (formerly GE Healthcare, now Cytiva) column and run at 25°C in SEC buffer (50 mM Tris pH 7.5, 150 mM Potassium acetate, 10 mM Magnesium acetate). The data was processed using the ASTRA software (Wyatt Technology).

#### Circular dichroism

CD experiments were recorded on a Jasco J-815 spectropolarimeter (Easton, MD) equipped with an automatic 6-position Peltier thermostated cell holder. The instrument was calibrated with 10-camphorsulphonic acid. Samples (65 μL) were prepared in PBS supplemented with CHAPS 7 mM and buffered at various pH (6.0, 7.0 or 8.0). Far-UV CD data were collected in the 182–270 nm range using a 0.1 mm pathlength cell (Quartz-Suprasil, Hellma UK Ltd) at 20.0 °C ± 0.1 °C. Spectra were acquired using a continuous scan rate of 50 nm/min and are presented as an average of 10 successive scans. The response time and the bandwidth were 1.0 s and 1 nm, respectively. The spectra were corrected by subtracting the solvent spectrum obtained under identical conditions.

#### CryoEM sample preparation

A 100μL BicD2 sample was thawed on ice and equilibrated with 5xPBS at either pH 8.0 or pH 7.0. The Superose^®^ 6 Increase column was equilibrated for 16h at a specific pH (in PBS buffer only) and the dimer fraction from the SEC taken for grid preparation. Cu/Rh 1.2/1.3 300mesh grids were plasma cleaned for 90s at 30% plasma power (80:20 argon:oxygen). The protein concentration was adjusted to 1μM and applied on the grid and plunge-frozen into liquid ethane in a Vitrobot Mark IV robot (FEI), maintained at 100% humidity and 10 °C.

#### CryoEM data collection and image analysis

Datasets for BicD2 at pH 8.0 and pH 7.0 were collected using a Glacios™ Cryo-TEM operating at 200keV with a K2 Summit direct electron detector (Gatan). Videos were collected in counting mode, with a final calibrated pixel size of 1.078 Å/pixel, 8s exposure, and total dose of ~ 54 e^-^/Å^2^.

SerialEM (Mastronarde, 2005) was used for automated data collection and the videos were processed using Relion3.0 (Zivanov et al., 2018). The micrographs were manually filtered giving each dataset ~700 micrographs. Particles were picked using the general model in TOPAZ (Bepler et al., 2019) and 2D classified twice. Each dataset yielded around 150k particles used for 2D classification. All data processing steps were done reference-free.

#### In vitro kinase assays and phosphorylation mapping

BicD2 or RanBP2_BBD_ were mixed with CDK1 at 1:100 molar ratio with 0.1mM MgATP and incubated for 30min at RT. Each sample had a negative control that did not contain MgATP. Each reaction mixture was loaded on SDS-PAGE and stained with pro-Q diamond stain to check for phosphorylated proteins. Phosphorylation sites were analysed with trypsin digest MS using an Orbitrap Elite and Acclaim Pepmap 100 column. The PSM values corresponded to the number of MS2 spectra which made it possible to identify peptides. In the case of BicD2 75% of the sequence were covered (PSM’s: 2087) and for RanBP2 85% of the sequence were covered (PSM’s: 1195).

#### AlphFold2 structure prediction

We used the whole CC2 and CC3 domain of a human BicD2 (residues 265-855) to model a homooligomer. We generated 5 models using the AlphaFold2 (Jumper et al., 2021) advanced notebook from Google Colab. We used MMseqs2 (Steinegger and Söding, 2017) (UniRef+Environmental) for MSA generation.

### QUANTIFICATION AND STATISTICAL ANALYSIS

No quantification or statistical analysis was carried out in this study.

## REFERENCES

Baffet, A.D., Hu, D.J., Vallee, R.B. (2015). Cdk1 Activates Pre-mitotic Nuclear Envelope Dynein Recruitment and Apical Nuclear Migration in Neural Stem Cells. Dev. Cell 33, 703–716. https://doi.org/10.1016/J.DEVCEL.2015.04.022.

Bepler, T., Morin, A., Rapp, M., Brasch, J., Shapiro, L., Noble, A.J., Berger, B., 2019. Positive-unlabeled convolutional neural networks for particle picking in cryo-electron micrographs. Nat. Methods 2019 1611 16, 1153–1160. https://doi.org/10.1038/s41592-019-0575-8.

Celestino, R., Henen, M.A., Gama, J.B., Carvalho, C., McCabe, M., Barbosa, D.J., Born, A., Nichols, P.J., Carvalho, A.X., Gassmann, R., Vögeli, B., 2019. A transient helix in the disordered region of dynein light intermediate chain links the motor to structurally diverse adaptors for cargo transport. PLoS Biol. 17. https://doi.org/10.1371/JOURNAL.PBIO.3000100.

Chambard, J.C., Pouyssegur, J., 1986. Intracellular pH controls growth factor-induced ribosomal protein S6 phosphorylation and protein synthesis in the G0→G1 transition of fibroblasts. Exp. Cell Res. 164, 282–294. https://doi.org/10.1016/0014-4827(86)90029-7.

Chowdhury, S., Ketcham, S.A., Schroer, T.A., Lander, G.C. (2015) Structural organization of the dynein-dynactin complex bound to microtubules. https://doi.org/10.1038/nsmb.2996.

d’Amico, E., Ud Din Ahmad, M., Cmentowski, V., Girbig, M., Müller, F., Wohlgemuth, S., Brockmeyer, A., Maffini, S., Janning, P., Vetter, I.R., Carter, A.P., Perrakis, A., Musacchio, A., 2022. Conformational transitions of the mitotic adaptor Spindly underlie its interaction with Dynein and Dynactin. bioRxiv 2022.02.02.478874. https://doi.org/10.1101/2022.02.02.478874.

Dodding, M.P., Way, M., 2011. Coupling viruses to dynein and kinesin-1. EMBO J. https://doi.org/10.1038/emboj.2011.283.

Fu, M. meng, Holzbaur, E.L.F., 2014. Integrated regulation of motor-driven organelle transport by scaffolding proteins. Trends Cell Biol. https://doi.org/10.1016/j.tcb.2014.05.002.

Gagliardi, L.J., Shain, D.H., 2013. Is intracellular pH a clock for mitosis? Theor. Biol. Med. Model. 10, 8. https://doi.org/10.1186/1742-4682-10-8.

Gama, J.B., Pereira, C., Simões, P.A., Celestino, R., Reis, R.M., Barbosa, D.J., Pires, H.R., Carvalho, C., Amorim, J., Carvalho, A.X., Cheerambathur, D.K., Gassmann, R., 2017. Molecular mechanism of dynein recruitment to kinetochores by the Rod-Zw10-Zwilch complex and Spindly. J. Cell Biol. 216, 943–960. https://doi.org/10.1083/JCB.201610108.

Gibson, J.M., Cui, H., Ali, M.Y., Zhao, X., Debler, E.W., Zhao, J., Trybus, K.M., Solmaz, S.R., Wang, C., 2021. Coil-to-Helix Transition at the Nup358-BicD2 Interface for Dynein Recruitment and Activation. bioRxiv 2021.05.06.443034. https://doi.org/10.1101/2021.05.06.443034.

Holland, P.M., Milne, A., Garka, K., Johnson, R.S., Willis, C., Sims, J.E., Rauch, C.T., Bird, T.A., Duke Virca, G. (2002). Purification, cloning, and characterization of Nek8, a novel NIMA-related kinase, and its candidate substrate Bicd2. J. Biol. Chem. 277, 16229–16240. https://doi.org/10.1074/JBC.M108662200.

Hoogenraad, C.C., Akhmanova, A., Howell, S.A., Dortland, B.R., De Zeeuw, C.I., Willemsen, R., Visser, P., Grosveld, F., Galjart, N. (2001). Mammalian golgi-associated Bicaudal-D2 functions in the dynein-dynactin pathway by interacting with these complexes. EMBO J. 20, 4041–4054. https://doi.org/10.1093/emboj/20.15.4041.

Hoogenraad, C.C., Wulf, P., Schiefermeier, N., Stepanova, T., Galjart, N., Small, J.V., Grosveld, F., De Zeeuw, C.I., Akhmanova, A. (2003). Bicaudal D induces selective dynein-mediated microtubule minus end-directed transport. EMBO J. 22, 6004–6015. https://doi.org/10.1093/emboj/cdg592.

Hu, D.J.K., Baffet, A.D., Nayak, T., Akhmanova, A., Doye, V., Vallee, R.B. (2013). Dynein recruitment to nuclear pores activates apical nuclear migration and mitotic entry in brain progenitor cells. Cell 154, 1300. https://doi.org/10.1016/J.CELL.2013.08.024.

Huynh, W., Vale, R.D. (2017). Disease-associated mutations in human BICD2 hyperactivate motility of dynein-dynactin. J. Cell Biol. 216, 3051–3060. https://doi.org/10.1083/JCB.201703201.

Indran, S. V, Ballestas, M.E., Britt, W.J. (2010). Bicaudal D1-dependent trafficking of human cytomegalovirus tegument protein pp150 in virus-infected cells. J. Virol. 84, 3162–3177. https://doi.org/10.1128/JVI.01776-09.

Jumper, J., Evans, R., Pritzel, A., Green, T., Figurnov, M., Ronneberger, O., Tunyasuvunakool, K., Bates, R., Žídek, A., Potapenko, A., Bridgland, A., Meyer, C., Kohl, S.A.A., Ballard, A.J., Cowie, A., Romera-Paredes, B., Nikolov, S., Jain, R., Adler, J., Back, T., Petersen, S., Reiman, D., Clancy, E., Zielinski, M., Steinegger, M., Pacholska, M., Berghammer, T., Bodenstein, S., Silver, D., Vinyals, O., Senior, A.W., Kavukcuoglu, K., Kohli, P., Hassabis, D. (2021). Highly accurate protein structure prediction with AlphaFold. Nat. 2021 1–7. https://doi.org/10.1038/s41586-021-03819-2.

Lee, I.-G., Olenick, M.A., Boczkowska, M., Franzini-Armstrong, C., Holzbaur, E.L.F., Dominguez, R. (2018). A conserved interaction of the dynein light intermediate chain with dynein-dynactin effectors necessary for processivity. Nat. Commun. 9. https://doi.org/10.1038/S41467-018-03412-8.

Liu, Y., Salter, H.K., Holding, A.N., Johnson, C.M., Stephens, E., Lukavsky, P.J., Walshaw, J., Bullock, S.L. (2013). Bicaudal-D uses a parallel, homodimeric coiled coil with heterotypic registry to coordinate recruitment of cargos to dynein. Genes Dev. 27, 1233–1246. https://doi.org/10.1101/GAD.212381.112.

Mastronarde, D.N. (2005). Automated electron microscope tomography using robust prediction of specimen movements. J. Struct. Biol. 152, 36–51. https://doi.org/10.1016/J.JSB.2005.07.007.

Matanis, T., Akhmanova, A., Wulf, P., Del Nery, E., Weide, T., Stepanova, T., Galjart, N., Grosveld, F., Goud, B., De Zeeuw, C.I., Barnekow, A., Hoogenraad, C.C. (2002). Bicaudal-D regulates COPI-independent Golgi-ER transport by recruiting the dynein-dynactin motor complex. Nat. Cell Biol. 4, 986–992. https://doi.org/10.1038/ncb891.

McClintock, M.A., Dix, C.I., Johnson, C.M., McLaughlin, S.H., Maizels, R.J., Hoang, H.T., Bullock, S.L. (2018). RNA-directed activation of cytoplasmic dynein-1 in reconstituted transport RNPs. Elife 7. https://doi.org/10.7554/ELIFE.36312.

McKenney, R.J., Huynh, W., Tanenbaum, M.E., Bhabha, G., Vale, R.D. (2014). Activation of cytoplasmic dynein motility by dynactin-cargo adapter complexes. Science 80. 345, 337–341. https://doi.org/10.1126/SCIENCE.1254198.

Noell, C.R., Loh, J.Y., Debler, E.W., Loftus, K.M., Cui, H., Russ, B.B., Zhang, K., Goyal, P., Solmaz, S.R. (2019). Role of Coiled-Coil Registry Shifts in the Activation of Human Bicaudal D2 for Dynein Recruitment upon Cargo Binding. J. Phys. Chem. Lett. 10, 4362–4367. https://doi.org/10.1021/acs.jpclett.9b01865.

Oates, E.C., Rossor, A.M., Hafezparast, M., Gonzalez, M., Speziani, F., MacArthur, D.G., Lek, M., Cottenie, E., Scoto, M., Foley, A.R., Hurles, M., Houlden, H., Greensmith, L., Auer-Grumbach, M., Pieber, T.R., Strom, T.M., Schule, R., Herrmann, D.N., Sowden, J.E., Acsadi, G., Menezes, M.P., Clarke, N.F., Züchner, S., Muntoni, F., North, K.N., Reilly, M.M. (2013). Mutations in BICD2 cause dominant congenital spinal muscular atrophy and hereditary spastic paraplegia. Am. J. Hum. Genet. 92, 965–973. https://doi.org/10.1016/J.AJHG.2013.04.018.

Olenick, M.A., Holzbaur, E.L.F. (2019). Dynein activators and adaptors at a glance. J. Cell Sci. 132. https://doi.org/10.1242/JCS.227132.

Paschal, B.M., Vallee, R.B. (1987). Retrograde transport by the microtubule-associated protein MAP 1C. Nature 330, 181–183. https://doi.org/10.1038/330181a0.

Peeters, K., Litvinenko, I., Asselberg, B., Almeida-Souza, L., Chamova, T., Geuens, T., Ydens, E., Zimoń, M., Irobi, J., Vriendt, E. De, Winter, V. De, Ooms, T., Timmerman, V., Tournev, I., Jordanova, A. (2013). Molecular defects in the motor adaptor BICD2 cause proximal spinal muscular atrophy with autosomal-dominant inheritance. Am. J. Hum. Genet. 92, 955–964. https://doi.org/10.1016/J.AJHG.2013.04.013.

Pouysségur, J., Franchi, A., L’Allemain, G., Paris, S. (1985). Cytoplasmic pH, a key determinant of growth factor-induced DNA synthesis in quiescent fibroblasts. FEBS Lett. 190, 115–119. https://doi.org/10.1016/0014-5793(85)80439-7.

Reck-Peterson, S.L., Redwine, W.B., Vale, R.D., Carter, A.P. (2018). The cytoplasmic dynein transport machinery and its many cargoes. Nat. Rev. Mol. Cell Biol. https://doi.org/10.1038/s41580-018-0004-3.

Rogerson, D.T., Sachdeva, A., Wang, K., Haq, T., Kazlauskaite, A., Hancock, S.M., Huguenin-Dezot, N., Muqit, M.M.K., Fry, A.M., Bayliss, R., Chin, J.W. (2015). Efficient genetic encoding of phosphoserine and its nonhydrolyzable analog. Nat. Chem. Biol. 2015 117 11, 496–503. https://doi.org/10.1038/nchembio.1823.

Schiavo, G., Greensmith, L., Hafezparast, M., Fisher, E.M.C. (2013). Cytoplasmic dynein heavy chain: The servant of many masters. Trends Neurosci. https://doi.org/10.1016/j.tins.2013.08.001.

Schlager, M.A., Hoang, H.T., Urnavicius, L., Bullock, S.L., Carter, A.P. (2014). In vitro reconstitution of a highly processive recombinant human dynein complex. EMBO J. 33, 1855–1868. https://doi.org/10.15252/EMBJ.201488792.

Schlager, M.A., Kapitein, L.C., Grigoriev, I., Burzynski, G.M., Wulf, P.S., Keijzer, N., Graaff, E. de, Fukuda, M., Shepherd, I.T., Akhmanova, A., Hoogenraad, C.C. (2010). Pericentrosomal targeting of Rab6 secretory vesicles by Bicaudal-D-related protein 1 (BICDR-1) regulates neuritogenesis. EMBO J. 29, 1637–1651. https://doi.org/10.1038/EMBOJ.2010.51.

Schroeder, C.M., Ostrem, J.M.L., Hertz, N.T., Vale, R.D. (2014). A Ras-like domain in the light intermediate chain bridges the dynein motor to a cargo-binding region. Elife 3, 1–22. https://doi.org/10.7554/ELIFE.03351.

Schroeder, C.M., Vale, R.D. (2016). Assembly and activation of dynein-dynactin by the cargo adaptor protein Hook3. J. Cell Biol. 214, 309–318. https://doi.org/10.1083/JCB.201604002.

Seale, J.W. (2006). The role of a conserved histidine–tyrosine interhelical interaction in the ion channel domain of δ-endotoxins from Bacillus thuringiensis. Proteins Struct. Funct. Bioinforma. 63, 385–390. https://doi.org/10.1002/PROT.20798.

Short, B., Preisinger, C., Schaletzky, J., Kopajtich, R., Barr, F.A. (2002). The Rab6 GTPase Regulates Recruitment of the Dynactin Complex to Golgi Membranes. Curr. Biol. 12, 1792–1795. https://doi.org/10.1016/S0960-9822(02)01221-6.

Sladewski, T.E., Billington, N., Ali, M.Y., Bookwalter, C.S., Lu, H., Krementsova, E.B., Schroer, T.A., Trybus, K.M. (2018). Recruitment of two dyneins to an mRNA-dependent bicaudal D transport complex. Elife 7. https://doi.org/10.7554/ELIFE.36306.

Splinter, D., Razafsky, D.S., Schlager, M.A., Serra-Marques, A., Grigoriev, I., Demmers, J., Keijzer, N., Jiang, K., Poser, I., Hyman, A.A., Hoogenraad, C.C., King, S.J., Akhmanova, A. (2012). BICD2, dynactin, and LIS1 cooperate in regulating dynein recruitment to cellular structures. https://doi.org/10.1091/mbc.e12-03-0210 23, 4226–4241.

Splinter, D., Tanenbaum, M.E., Lindqvist, A., Jaarsma, D., Flotho, A., Yu, K. Lou, Grigoriev, I., Engelsma, D., Haasdijk, E.D., Keijzer, N., Demmers, J., Fornerod, M., Melchior, F., Hoogenraad, C.C., Medema, R.H., Akhmanova, A. (2010). Bicaudal D2, dynein, and kinesin-1 associate with nuclear pore complexes and regulate centrosome and nuclear positioning during mitotic entry. PLoS Biol. 8. https://doi.org/10.1371/journal.pbio.1000350.

Steinegger, M., Söding, J. (2017). MMseqs2 enables sensitive protein sequence searching for the analysis of massive data sets. Nat. Biotechnol. 2017 3511 35, 1026–1028. https://doi.org/10.1038/nbt.3988.

Stuurman, N., Häner, M., Sasse, B., Hübner, W., Suter, B., Aebi, U. (1999). Interactions between coiled-coil proteins: Drosophila lamin Dm0 binds to the Bicaudal-D protein. Eur. J. Cell Biol. 78, 278–287. https://doi.org/10.1016/S0171-9335(99)80061-2.

Terawaki, S.I., Yoshikane, A., Higuchi, Y., Wakamatsu, K. (2015). Structural basis for cargo binding and autoinhibition of Bicaudal-D1 by a parallel coiled-coil with homotypic registry. Biochem. Biophys. Res. Commun. 460, 451–456. https://doi.org/10.1016/j.bbrc.2015.03.054.

Torisawa, T., Ichikawa, M., Furuta, A., Saito, K., Oiwa, K., Kojima, H., Toyoshima, Y.Y., Furuta, K. (2014). Autoinhibition and cooperative activation mechanisms of cytoplasmic dynein. Nat. Cell Biol. 2014 1611 16, 1118–1124. https://doi.org/10.1038/ncb3048.

Urnavicius, L., Zhang, K., Diamant, A.G., Motz, C., Schlager, M.A., Yu, M., Patel, N.A., Robinson, C. V, Carter, A.P. (2015). The structure of the dynactin complex and its interaction with dynein. Science 347, 1441–1446. https://doi.org/10.1126/SCIENCE.AAA4080.

Van Damme, P., Lasa, M., Polevoda, B., Gazquez, C., Elosegui-Artola, A., Kim, D.S., De Juan-Pardo, E., Demeyer, K., Hole, K., Larrea, E., Timmerman, E., Prieto, J., Arnesen, T., Sherman, F., Gevaert, K., Aldabe, R. (2012). N-terminal acetylome analyses and functional insights of the N-terminal acetyltransferase NatB. Proc. Natl. Acad. Sci. U. S. A. 109, 12449–12454. https://doi.org/10.1073/PNAS.1210303109/-/DCSUPPLEMENTAL.

Veld, P.J.H. in ‘t, Wohlgemuth, S., Koerner, C., Mueller, F., Janning, P., Musacchio, A. (2022). Reconstitution and use of highly active human CDK1:Cyclin-B:CKS1 complexes. Protein Sci. 31, 528–537. https://onlinelibrary.wiley.com/doi/10.1002/pro.423.

Wanschers, B.F.J., van de Vorstenbosch, R., Schlager, M.A., Splinter, D., Akhmanova, A., Hoogenraad, C.C., Wieringa, B., Fransen, J.A.M. (2007). A role for the Rab6B Bicaudal–D1 interaction in retrograde transport in neuronal cells. Exp. Cell Res. 313, 3408–3420. https://doi.org/10.1016/J.YEXCR.2007.05.032.

Wharton, R.P., Struhl, G. (1989). Structure of the Drosophila BicaudalD protein and its role in localizing the posterior determinant nanos. Cell 59, 881–892. https://doi.org/10.1016/0092-8674(89)90611-9.

Wilson, M.H., Holzbaur, E.L.F. (2012). Opposing microtubule motors drive robust nuclear dynamics in developing muscle cells. J. Cell Sci. 125, 4158–4169. https://doi.org/10.1242/jcs.108688.

Zhang, K., Foster, H.E., Rondelet, A., Lacey, S.E., Bahi-Buisson, N., Bird, A.W., Carter, A.P. (2017). Cryo-EM Reveals How Human Cytoplasmic Dynein Is Auto-inhibited and Activated. Cell 169, 1303–1314.e18. https://doi.org/10.1016/J.CELL.2017.05.025.

Zivanov, J., Nakane, T., Forsberg, B.O., Kimanius, D., Hagen, W.J.H., Lindahl, E., Scheres, S.H.W. (2018). New tools for automated high-resolution cryo-EM structure determination in RELION-3. Elife 7. https://doi.org/10.7554/ELIFE.42166.

