## Supplementary material for "*In vitro* characterization of the full-length human dynein-1 cargo adaptor BicD2": Key resource table

| REAGENT or RESOURCE | SOURCE | IDENTIFIER |
|---|---|---|
| Bacterial and insect cell strains strains | | |
| *E. coli* BL21(DE3) | New England Biolabs | C2527H |
| Sf9 insect cells | Sigma-Aldrich | 71104-M |
| Oligonucleotides | | |
| To linearize pNHD vector, forward primer: ATGTATAATCTCCTTCTTAA | Eurofins | N/A |
| To linearize pNHD vector, reverse primer: TAATGAGGGGTACCCTTGGG | Eurofins | N/A |
| To subclone into pNHD vector, forward primer: TCATCAGCGTGGTCGTGAAGCGATT | Eurofins | N/A |
| To subclone into pNHD vector, reverse primer: CTCAACGAGCTGGACGCGGATGA | Eurofins | N/A |
| To amplify RanBP2 gene from pET vector, forward primer: ATGACTGAAGATTCCGATGATATCC | Eurofins | N/A |
| To amplify RanBP2 gene from pET vector, reverse primer: TTAGTGATGATGATGATGATGGCTG | Eurofins | N/A |
| To attach RanBP2 gene overlap to pNHD, reverse primer: CTTCAGTCATATGTATAATCTCCTTCTTAA | Eurofins | N/A |
| To attach RanBP2 gene overlap to pNHD, forward primer: CATCACTAATAATGAGGGGTACCCTTGGGA | Eurofins | N/A |
| To attach BicD2 gene overlap to pNHD, reverse primer: AGACCACATATGTATAATCTCCTTCTTAA | Eurofins | N/A |
| To attach BicD2 gene overlap to pNHD, forward primer: TGCGACTAATAATGAGGGGTACCCTTGGGA | Eurofins | N/A |
| To introduce Y538A and H539A mutations in BicD2, forward primer: CATGAAGACGGTGACGCGGCGGAGGTTGA CATCAAC | Eurofins | N/A |
| To introduce Y538A and H539A mutations in BicD2, forward primer: GTTGATGTCAACCTCCGCCGCGTCACCGTC TTCATG | Eurofins | N/A |
| To introduce H640A mutation in BicD2, forward primer: CGTATCGCGAGCCAAGCGCTGGGTCCAGC CGTT | Eurofins | N/A |
| To introduce H640A mutation in BicD2, reverse primer: AACGGCTGGACCCAGCGCTTGGCTCGCGAT ACG | Eurofins | N/A |
| Recombinant DNA | | |
| pNHD plasmid | Rogerson et al., 2015 | N/A |
| Human RanBP2 in pET42 vector | Epoch Life Science | N/A |
| Human BicD2 in pET42 vector | Epoch Life Science | N/A |
| Software and algorithms | | |
| AlphaFold2 | Jumper et al., 2021 | https://github.com/deepmind/alphafold |

| | | |
|---|---|---|
| Relion | Zivanov et al., 2018 | https://github.com/3dem/relion |
