## Supplementary figures for "*In vitro* characterization of the full-length human dynein-1 cargo adaptor BicD2"

A

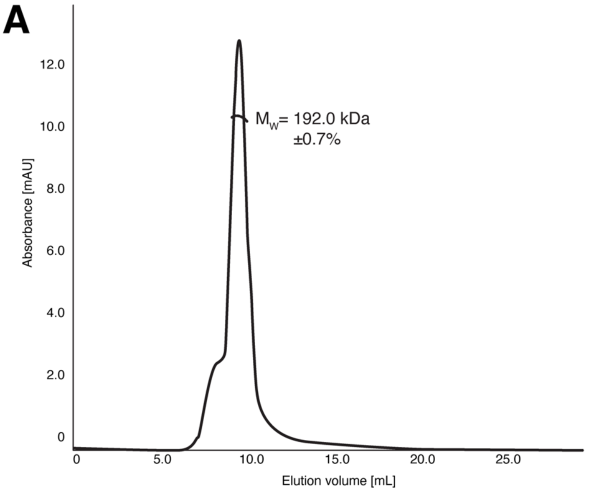


B

| | Charge, z | Net charge at pH7.4 | Isoelectric point |
|---|---|---|---|
| $BicD2_{FL}$ | -16.61 (pH6.0) | -31.92 | 5.37 |
| | -29.52 (pH7.0) | | |
| | -35.66 (pH8.0) | | |

C

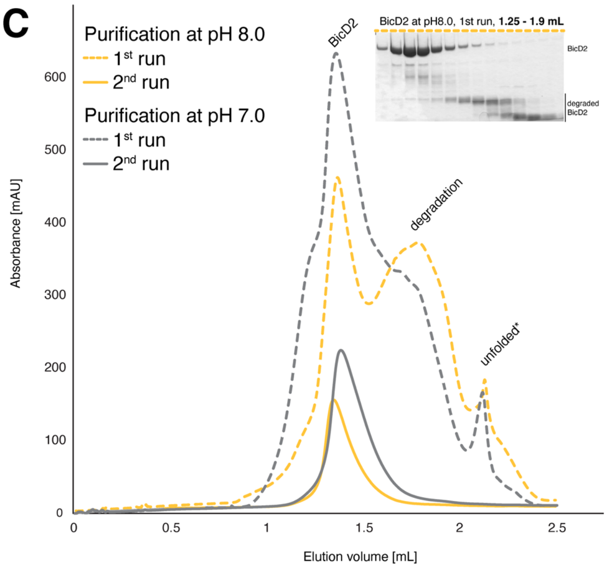


D

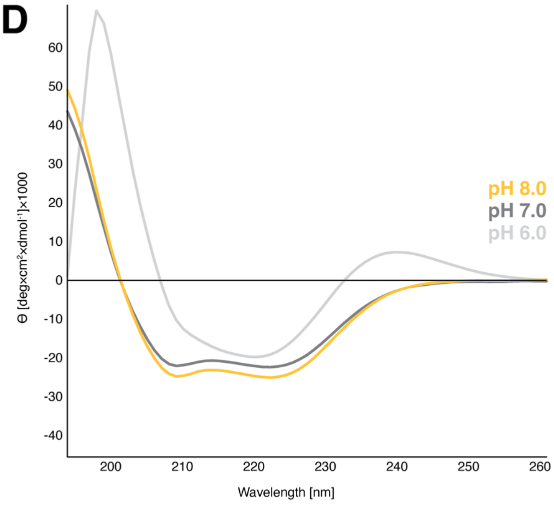


E

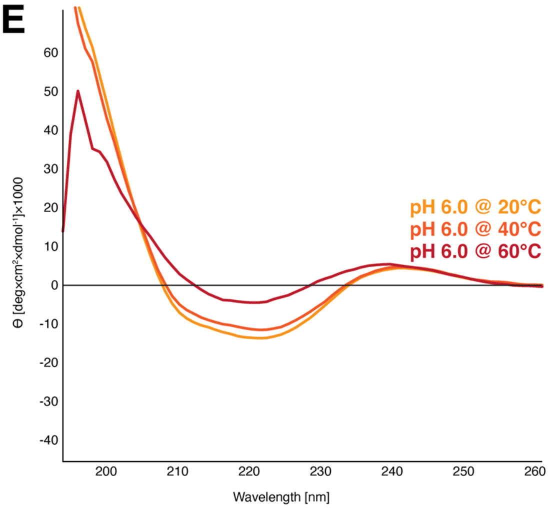

**Supplementary figure 1. Additional biophysical characterization of human full-length BicD2, Related to Figure 2.** (A) The SEC-MALS profile of pre-purified BicD2. The value of 192kDa ± 0.7% is in a good correlation with a theoretical BicD2 dimer weight of 186kDa considering the residual CHAPS detergent present in the sample (CHAPS micelle ≈6kDa). (B) Table with BicD2 charge values at specific pH values. To generate this data we used the 'Prot pi' bioinformatic tool box for the calculation and simulation of physico-chemical parameters (Release: 2.2.29.150). (C) SEC profile of full-length BicD2. The grey chromatogram represents the sample ran at pH 7.0, and the yellow chromatogram is the sample ran at pH 8.0. The dashed lines represent SEC profiles directly after affinity purification. The presence of degradation products was confirmed by SDS-PAGE (shown in the figure) and MS analysis. Peak fractions from these runs were pooled and re-run (solid lines) to obtain homogenous BicD2 fractions that are free of degradation products. In both cases an identical protein amount was loaded on to the Superose® 6 Increase column. (D) CD spectrum of the full-length BicD2 at pH values of 6.0, 7.0 and 8.0. All three runs were done at 20°C. (E) CD spectrum of full-length BicD2 at pH 6.0 at three temperatures (20, 40, and 60°C). Prior to the experiment each BicD2 sample was purified twice by SEC in PBS buffer at the respective pH.

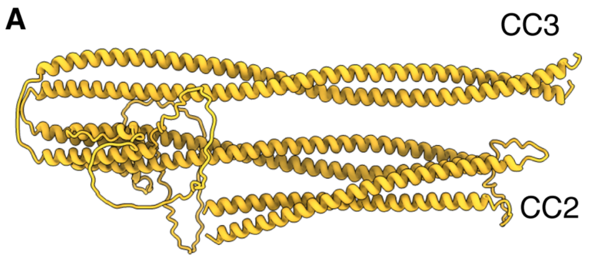


AF2 model of BicD2$_{CC2\text{-}CC3}$

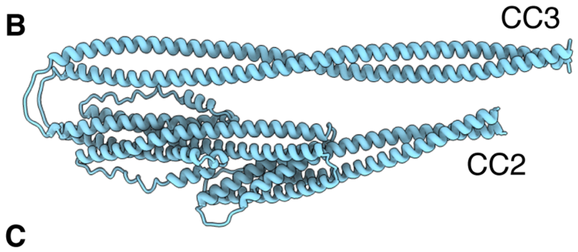


AF2 model of BicD$_{CC2\text{-}CC3}$

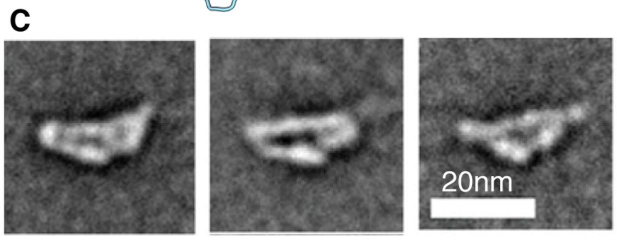


Negative stain class averages of a full-length BicD from Sladewski *et al.*, 2018

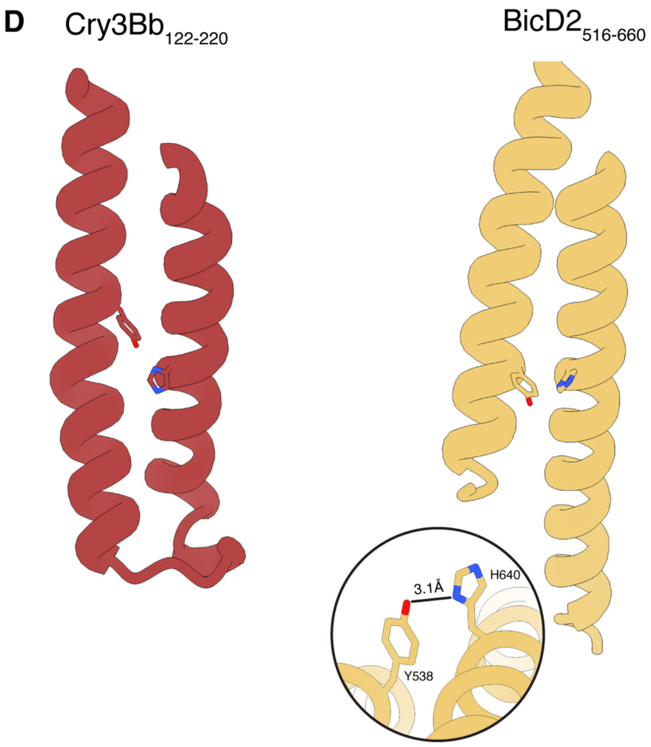


```
212  QVFGEEWGYSSEDVAEFYHRQLKLTQQYTDH-CVNWYNVGLNGLRGST 259
521  SVAQDELVTFSEELANLYHHVCMCNNETPNRVMLDYYREGQGGAGRTS 568
     .*  :*    **::*::**:    .:: .::  :::*. * .*   ::
```

**Supplementary figure 2. Validation and analysis of the AF2 generated BicD2$_{CC2-CC3}$ model, Related to Figure 3.** Comparison of the (A) AF2-predicted human BicD2$_{CC2-CC3}$ structure, (B) the AF2-predicted fruit fly BicD$_{CC2-CC3}$ structure, and (C) negative stain class-averages of the full-length BicD from Sladewski *et al*., 2018. (D) Structure comparison of the predicted structure of the BicD2 H3-H4-H5 α-helical bundle fragment and the analogous region in Cry3Bb endotoxin (PDB ID: 1CIY) and sequence alignment of the tyrosine containing helix. Loops removed for clarity reasons. The circle shows the distance from Y538 and H640 in the predicted BicD2 structure.

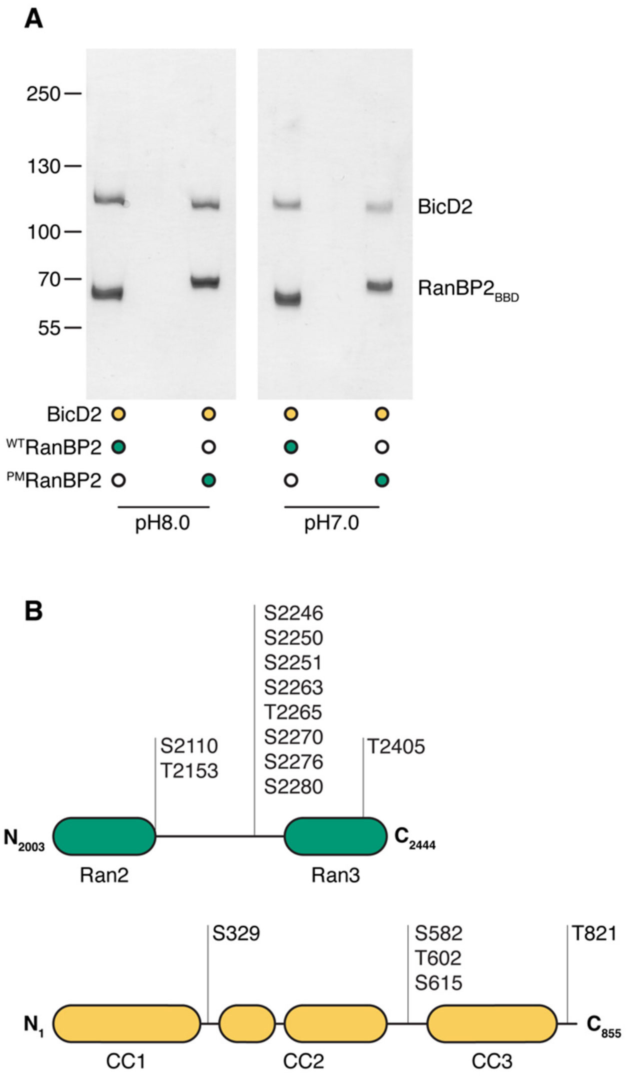


**Supplementary figure 3. BicD2-RanBP2$_{BBD}$ interaction and phosphorylation analysis of RanBP2$_{BBD}$ and BicD2, Related to Figure 4.** (A) Double pull-down of the BicD2-RanBP2$_{BBD}$ complex. The first pull-down was done on Strep-tagged BicD2 followed by 6xHis-tagged RanBP2. The complex was pulled with either RanBP2$_{BBD}$ (RanBP2$_{BBD}$-WT) or the phosphomimetic mutant of RanBP2$_{BBD}$ (RanBP2$_{BBD}$-PM) at two different pH values. (B) CDK1-CyclinB phosphorylation profile of BicD2 and RanBP2. RanBP2$_{BBD}$ contains multiple phosphorylation sites within the structurally disordered region between the Ran2 and Ran3 domains, which is also the BicD2 binding site. BicD2 contains five S/T sites within its sequence in structurally disordered regions.

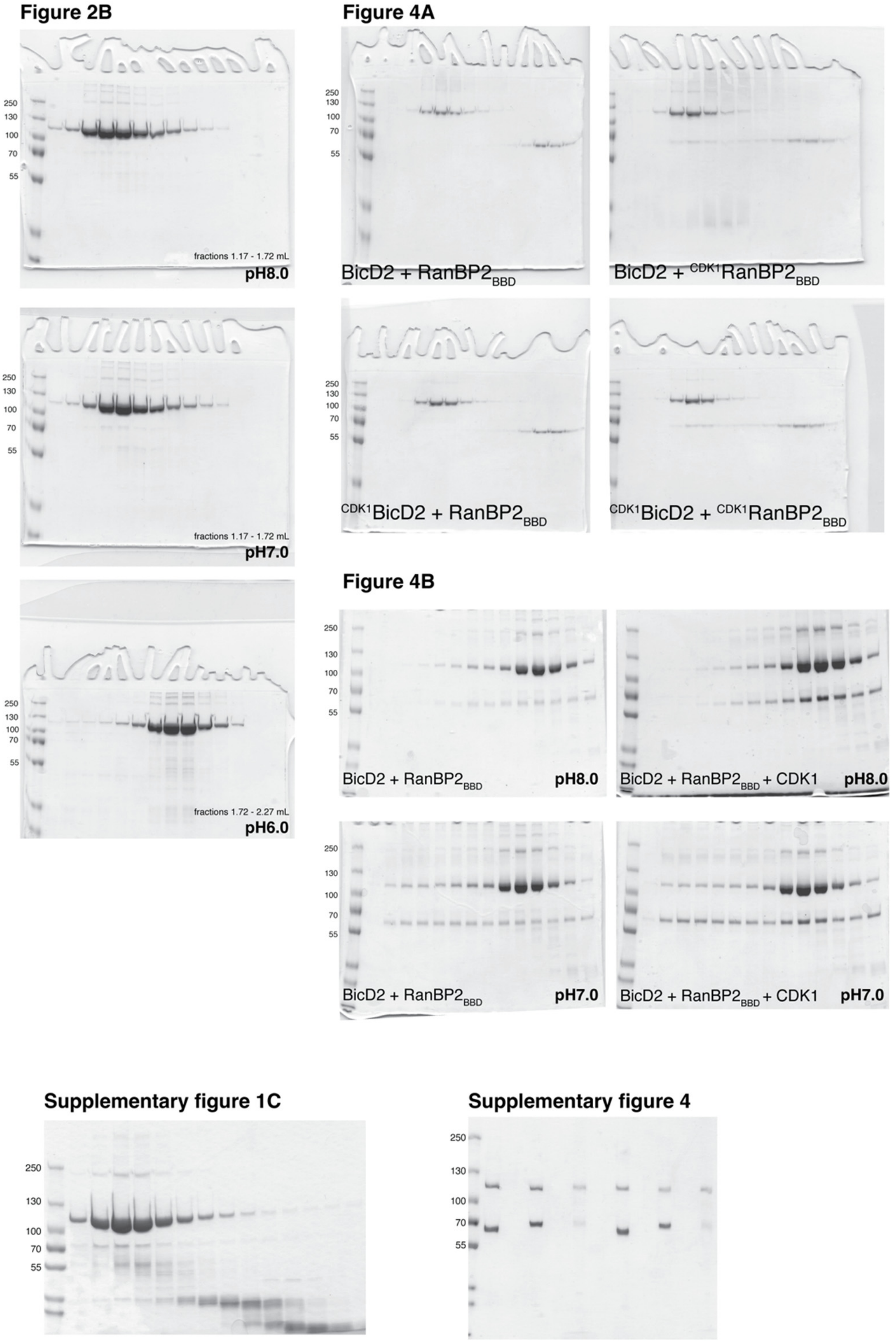


**Supplementary figure 4. Original SDS-PAGE gels presented in this work, Related to Figures 2 and 4 as well as Supplementary figures 1 and 4.** The identities of BicD2 and RanBP2 were confirmed by MS.
